# Nongenic cancer-risk SNPs affect oncogenes, tumor suppressor genes, and immune function

**DOI:** 10.1101/507236

**Authors:** M. Fagny, J. Platig, M.L. Kuijjer, X. Lin, J. Quackenbush

## Abstract

Genome-wide associations studies (GWASes) have identified many germline genetic variants that are associated with an increased risk of developing cancer. However, how these single nucleotide polymorphisms (SNPs) alter biological function in a way that increases cancer risk is still largely unknown. We used a systems biology approach to analyze the regulatory role and functional associations of cancer-risk SNPs in thirteen distinct tissues. Using data from the Genotype-Tissue Expression (GTEx) project, we performed an expression quantitative trait locus (eQTL) analysis, keeping both *cis*- and *trans*-eQTLs, and representing those significant associations as edges in tissue-specific eQTL bipartite networks. We find that each network is organized into highly modular communities that group sets of SNPs together with functionally-related collections of genes. We mapped cancer-risk SNPs to each tissue-specific eQTL network. Although we find in each tissue that cancer-risk SNPs are distributed across the network, they are not uniformly distributed. Rather they are significantly over-represented in a small number of communities. This includes communities enriched for immune response processes as well as communities representing tissue-specific functions. Moreover, cancer-risk SNPs are over-represented in the central “cores” of communities, meaning they are more likely to influence the expression of many genes within the same community, thus affecting biological processes. And finally, we find that cancer-risk SNPs preferentially target oncogenes and tumor suppressor genes, suggesting non-genic mutations may still alter the effects of these key cancer-associated genes. This bipartite eQTL network approach provides a new way of understanding genetic effects on cancer risk and provides a biological context for interpreting the results of GWAS cancer studies.

## Introduction

Cancers often result from somatic mutations in oncogenes and tumor suppressors, which frequently arise due to environmental exposures such as UV light, tobacco, smoke, or chemicals. Hereditary cancers are representing between 5 and 10% of all cancers and are characterized by a family history of the disease, a younger than usual age of onset, and a higher likelihood of primary cancers in multiple organs. They are often associated with germline alterations in these same classes of genes, called oncogenes or tumor suppressor genes. However, beyond these obvious candidate cancer drivers, it is widely recognized that other genetic factors play a role in development and cancer progression. Genome Wide Association Studies (GWASes) have identified germline single nucleotide polymorphisms (SNPs) that are associated with altered cancer risk (“cancer-risk SNPs”), including *BRCA1* and *BRCA2* gene mutations known to increase risk of breast and ovarian cancers. However, many SNPs identified through GWASes fall into non-genic regions, making it difficult to interpret their biological role in disease development, progression, and response to therapy.

The population frequency of a germline cancer-risk SNP is generally anti-correlated with its effect, calculated as the relative risk between people who carry the mutation and those who do not [1]. Although the functions of the small number of rare variants with strong effects are well-studied, little is known about the functions of the more common risk variants with small effects that are present at intermediate frequency in the general population. Among the SNPs in the GWAS catalog that pass the genome-wide significance bar for association with an elevated risk for one or more cancers, most have an odds ratio less than 1.3, and most fall outside of genes (in “non-genic” regions), suggesting they may play a role in the regulation of gene expression [2, 1].

Expression quantitative traits locus (eQTL) analysis tests for associations between the genotype at a SNP locus and expression levels of a gene, and an eQTL association can provide evidence for a SNP’s regulatory role. GWASes have identified germline SNPs that are linked to cancer risk and a number of studies have found that these SNPs influence gene expression levels [3, 4, 5]. However, no study has systematically investigated the biological characteristics and functional impact of regulatory germline “cancer risk” SNPs in the general population.

This gap in our understanding of cancer-risk SNPs may be due to their inherent characteristics. In addition to their small effect on the macroscopic phenotype (developing cancer), cancer-risk SNPs also usually have small effects on the expression of individual genes. Moreover, because many genes exhibit tissue-specific expression, it is difficult to characterize the regulatory role of cancer-risk SNPs that target genes not expressed in the most frequently studied tissues, such as whole blood. Finally, because the transformation of a healthy cell into a cancer cell is associated with many genomic and transcriptomic changes, we cannot use the studies of tumor cells to investigate the effect of the regulatory cancer-risk SNPs on pre-tumor cells.

However, new data sets can help shed light on the role of cancer-risk SNPs. Large-scale studies, such as the Genotype-Tissue Expression (GTEx) project, provide genomic and transcriptomic data from hundreds of individuals and dozens of non-diseased tissues [6], thus allowing the effects of cancer-risk SNPs to be assessed in multiple tissues, including those in which their effects are most relevant.

In this study, we used a system biology approach to characterize the regulatory role of germline cancer-risk SNPs in thirteen different tissues (Table S2) using data from the GTEx project v6.0. In each tissue, we performed an eQTL analysis and represented both *cis*- and *trans*-eQTLs using a bipartite network. We then mapped both germline cancer-risk SNPs and the oncogenes and tumor suppressor genes to these networks and used the properties of the networks to identify the biological functions and pathways that cancer-risk SNPs affect.

We find that although cancer-risk SNPs are distributed across the network, they are enriched in a small number of communities associated with immune response and recognition of pathogens, underscoring the importance of immune processes in cancer. In particular, cancer-risk SNPs preferentially map to communities enriched for genes belonging to the Major Histocompatibility Complex (MHC) indicating a potentially greater role for immune processes in cancer risk than might have been expected. We also find that cancer-risk SNPs are overrepresented among local community hubs (“core SNPs”), regulating multiple genes involved in the same biological function both in *cis* and in *trans*. Finally, we find that cancer-risk SNPs are preferentially located in the promoters of oncogenes and tumor suppressor genes and are more likely than expected by chance to influence the expression level of these cancer-related genes. This analysis demonstrates the power of using tissue-specific bipartite eQTL networks as a framework to investigate how germline SNPs can act coordinately to deregulate the expression of biological functions and lead to an increased risk to develop cancer.

## Results

### Cancer risk SNPs are located in non-coding regions

We defined a set of cancer-risk SNPs based on the NHGRI-EBI GWAS catalog (accession date: 2017-04-24); we extracted a set of 874 SNPs from 565 independent linkage disequilibrium (LD) blocks associated (at genome-wide significance *p* ≤ 5 × 10^−8^) with 130 unique traits and diseases terms related to cancers, representing 41 cancer types (see Supplementary Table S1). Most of the cancer-risk SNPs were associated with only one cancer type; only 6% were associated with two or more cancers, and only 2% with more than three cancers. In contrast, most cancer types (82%) were associated with multiple independent SNPs, with number of associated independent SNPs ranging between 1 (B cell non-Hodgkin lymphoma, cardiac gastric cancer, chronic myeloid leukemia, meningioma, non-melanoma skin cancer, small intestine neuroendocrine tumor, and sporadic pituitary adenoma) and 95 (prostate cancer).

When examining the genomic location of cancer-risk SNPs, we found that their individual effect on the risk of developing cancer was also generally small with over 99% of cancer-risk SNPs having an odds ratio under 3. To our surprise, we found that only 9.7% of cancer-risk SNPs were exonic or splice variant SNPs. The lack of a clear known biological function based on SNP location suggests that many of the remaining 91.3% may play a regulatory role.

### Cancer-risk SNPs regulate cancer-related biological functions

To characterize the biological functions of this large number of small-effect, regulatory, cancer-risk SNPs, we performed a systems-based eQTL analysis of genotyping and RNA-Seq data from GTEx v6.0. After filtering and normalizing the GTEx data, and eliminating tissues for which there were fewer than 200 samples, we were left with gene expression and genotype data for thirteen tissues (twelve primary tissues and one cell line, see Supplementary Table S2). We used MatrixeQTL [7], correcting for reported sex, age, ethnic background and the top three genotype principal components, to compute eQTLs in cis and *trans,* within a +/-1Mb window around the genes (see Methods).

For each of the thirteen tissues, we represented the significant *cis* - and *trans*-eQTL results at a FDR threshold less than or equal to 0.2 as a bipartite network, where nodes are either SNPs or genes and edges are significant associations between SNPs and genes[8],[9]. We obtained thirteen tissue-specific networks containing between 57,641 (ATA—aorta) and 431,036 (THY—thyroid) SNPs (median across all thirteen tissues = 198,226), corresponding to between 3,550 and 34,016 LD blocks (median = 15,514), and between 1,090 and 10,003 genes (median = 4,820). We used R condor package [8] in each of the thirteen eQTL networks to identify communities, defined as groups of SNPs and genes more densely connected to each other than would be expected by chance (see Methods). The modularity of these networks ranges from 0.83 to 0.97 (median = 0.95), indicating that they are highly structured, with SNPs and genes grouped in well-defined communities; in the thirteen tissues we found between 29 and 177 (median = 124) communities. We then functionally annotated those communities by testing for over-representation of genes annotated to Gene Ontology (GO) biological processes [10] (Supplementary Table S3).

We mapped the cancer-risk SNPs to the eQTL network for each of the thirteen tissues and found that cancer-risk SNPs appear in communities associated with a wide range of biological processes. Depending on the tissue, between 21% (heart left ventricle) and 49% (lung) of communities contain at least 1 cancer-risk SNPs. However, most communities contain only one or two cancer-risk SNPs (Table 1 and Figure 1A). A complete list of the cancer-risk snps mapping to the communities in each 13 tissues and their corresponding Gene Ontology Biological processes is provided in Supplementary Table S4). Using Fisher’s exact test, we identified 2 to 8 (median = 4) communities in each tissue that were enriched for SNPs associated with increased risk of any cancer (independent of tissue), and only a very small number that were enriched for cancer-risk SNPs associated with one particular type of cancer (Table 1). The details about enrichments, odds ratios, and p-values for each cancer type, each community, and each tissue are given in Supplementary Table S5.

**Table 1:** Communities enriched in cancer-risk SNPs

| **Tissue** | **Abbrev** | **# Communities** | **# With cancer risk SNPs** | **# Enriched in cancer risk SNPs<sup>a</sup>** | **# Enriched in 1+ cancer type<sup>b</sup>** |
| --- | --- | --- | --- | --- | --- |
| Adipose subcutaneous | ADS | 82 | 39 | 4 | 1 |
| Aorta | ATA | 29 | 13 | 2 | 2 |
| Artery tibial | ATT | 95 | 40 | 2 | 0 |
| Fibroblast | FIB | 156 | 54 | 2 | 3 |
| Esophagus mucosa | EMC | 147 | 45 | 4 | 2 |
| Esophagus muscularis | EMS | 143 | 58 | 2 | 1 |
| Heart left ventricle | HRV | 124 | 26 | 2 | 2 |
| Lung | LNG | 35 | 17 | 4 | 1 |
| Skeletal muscle | SMU | 86 | 33 | 3 | 1 |
| Tibial nerve | TNV | 152 | 64 | 8 | 1 |
| Skin | SKN | 163 | 71 | 4 | 3 |
| Thyroid | THY | 177 | 77 | 6 | 2 |
| Whole blood | WBL | 66 | 32 | 5 | 2 |

<sup>a</sup>In this column, all cancer risk SNPs across all cancer types were pooled together under a "cancer risk" label before the enrichment analysis was performed.
<sup>b</sup>In this column, each cancer type was analysed separately in each community. One community can be enriched for risk SNPs for multiple cancers

**Figure 1:**
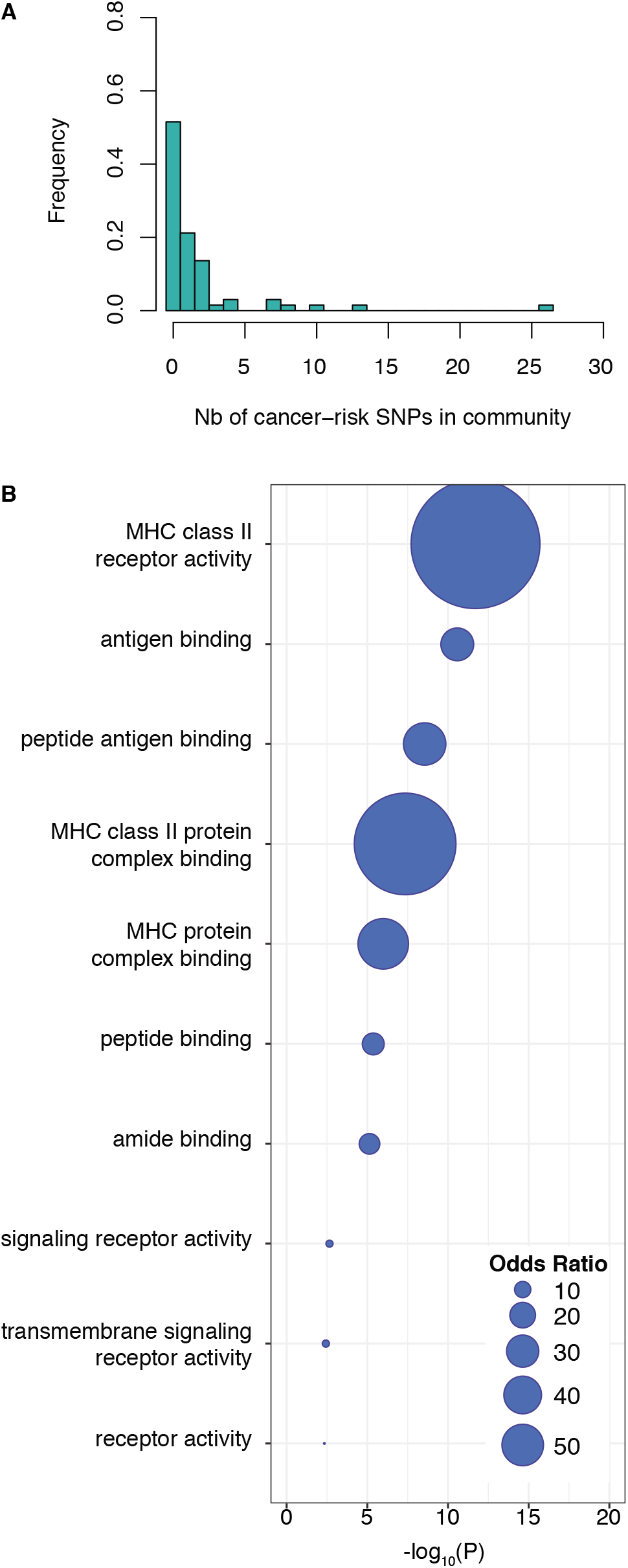
Cancer-risk SNPs are distributed across the network communities and functional roles. **A.** Distribution of the number of cancer-risk SNPs in each community in whole blood. **B.** Gene Ontology Term enrichment for communities in whole blood that are also enriched for cancer-risk SNPs.

We explored the functional consequences of cancer-risk SNPs by testing whether communities enriched for these SNPs were also enriched for genes annotated to GO biological process terms. Across all tissues except tibial artery (ATA), we found that communities with increased representation of cancer-risk SNPs contain genes enriched in functions linked to immunity, mainly genes belonging to the Major Histocompatibility Complex (MHC) class I and II families, and that the majority of these immune-related genes were cis-eQTLs with cancer-risk SNPs. An example of the Gene Ontology enrichment of this shared community in whole blood is presented Figure 1B. Other communities were enriched in non-specific biological processes like RNA metabolic processes and DNA binding. Only two of the tissue-specific networks presented a community enriched in both cancer-risk SNPs and tissue-specific biological pathways: epithelium development in skin and cell-cell adhesion in fibroblasts (Supplementary Table S3).

### Cancer risk SNPs are core SNPs in their communities

As shown previously, the communities in eQTL networks have a characteristic structure, with local hubs, or “core SNPs,” central within their communities. Disease-associated SNPs found through GWAS map not only to communities with relevant biological functions, but also to the cores of those communities[8, 9]. As a measure of SNP centrality, we define a “core score” equal to the relative modularity contributed by a SNP to the overall modularity of its community (see Eq. 2 in Methods). We calculated core scores for all SNPs in the network and compared the core score distribution for cancer-risk SNPs and SNPs not associated with any trait or disease in GWAS. We found that cancer-risk SNPs were enriched for higher core-scores (Figure 2A). This result is consistent across tissues (Supplementary Figure S2), indicating that germline cancer-risk SNPs, being central to their communities, affect the expression of many genes involved in coherent biological processes related to cancer development and progression.

**Figure 2:**
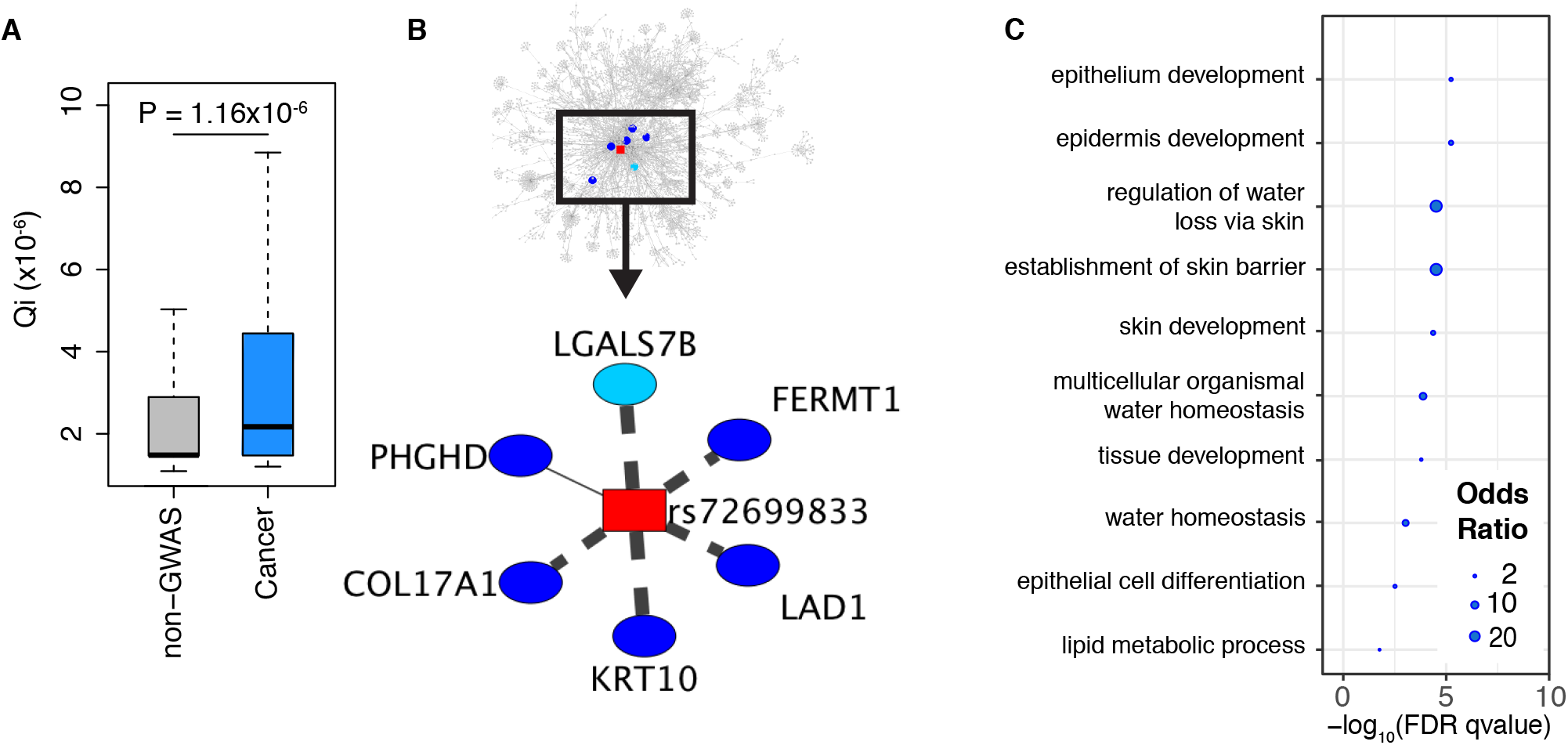
Network properties of GWAS cancer-risk SNPs. **A.** Distribution of core scores for SNPs associated with increased cancer risk in skin by GWAS (in blue) and other skin SNPs (in gray). P-values were obtained using a likelihood ratio test and pruning for SNPs in linkage disequilibrium. Distributions for all tissue-specific networks are in Supplementary Figure S2. **B.** An example of a SNP with high core-score: rs72699833, in LD with rs11249433, a SNP associated with a higher risk to develop breast cancer. This SNP belongs to community 147 (top panel) which is enriched for breast cancer-risk SNPs and is associated with multiple genes involved in epithelium development. LGALS7B is represented here but belongs to another community (107). Details on the associations are provided in Supplementary Table S6. Dashed lines indicate association in *trans*, full line in *cis*. The thickness of the lines correspond to the strength of the association. **C.** Enrichment in Gene Ontology Terms for community 147 in skin.

For example, SNP rs72699833 is a core SNP in skin community 147. This SNP is in LD with rs11249433, which has been associated with an increased risk of breast cancer (Figure 2B). When examining skin community 147, we find it to be enriched for SNPs associated with breast cancer (Supplementary Table S5) and for genes involved in epithelium development (Supplementary Table S3 and Figure 2C); as breast cancer is a epithelial cancer, the association with skin is not surprising. SNP rs72699833 is located on chromosome 1 and is associated in *cis* to *PHGDH,* a gene involved in the metabolism of serine and is over-expressed in some subtypes of breast, cervical, colorectal and non-small cells lung cancer and generally associated with a poorer outcome [11, 12, 13, 14].

In addition, rs72699833 is associated through our eQTL analysis in *trans* with five other genes: *LAD1* on chromosome 1, *COL17A1* on chromosome 10, *KRT10* on chromosome 17, *LGALS7B* on chromosome 19 and *FERMT1* on chromosome 20 (Supplementary Table S6). All of these genes are involved in epithelium development and in particular with extra-cellular matrix (ECM) secretion and cell-ECM interactions. Most of these genes have been shown to be dysregulated in breast cancer or during epithelial-mesenchymal transition. Indeed, *LAD1* has been associated with aggressive breast tumors [15], *COL17A1* is under-expressed in breast cancer and over-expressed in head and neck squamous cell carcinoma, lung squamous cell carcinoma, and lung adenocarcinoma [16] and *FERMT1* is a know mediator of epithelial–mesenchymal transition in colon cancer [17].

### Cancer risk SNPs preferentially target cancer genes

We expected that cancer-risk SNPs might be preferentially associated with genes known to be involved in cancer development and progression. We assembled a catalog of oncogenes and tumor suppressor genes (“cancer genes”) using databases that included the Network Gene Cancer version 5.0 [18] and the COSMIC [19] census (see Methods and Supplementary Table S7).

We tested whether cancer-risk SNPs are more frequently associated with cancer genes than other SNPs based on the eQTL networks. We mapped cancer-risk SNPs to the giant connected component of each of the thirteen tissue-specific eQTL networks. We then compared the number of cancer genes targeted by cancer-risk SNPs and others SNPs, taking into account linkage disequilibrium and global degree distribution (the total number of genes to which they were associated; see Methods). We showed that cancer-risk SNPs were indeed more likely to target cancer genes than expected by chance (*p* < 10^−6^ based on 1,000,000 resamplings) when studying all networks together (Figure 3B); similar results were found in each tissue-specific network (Supplementary Figure S1).

**Figure 3.**
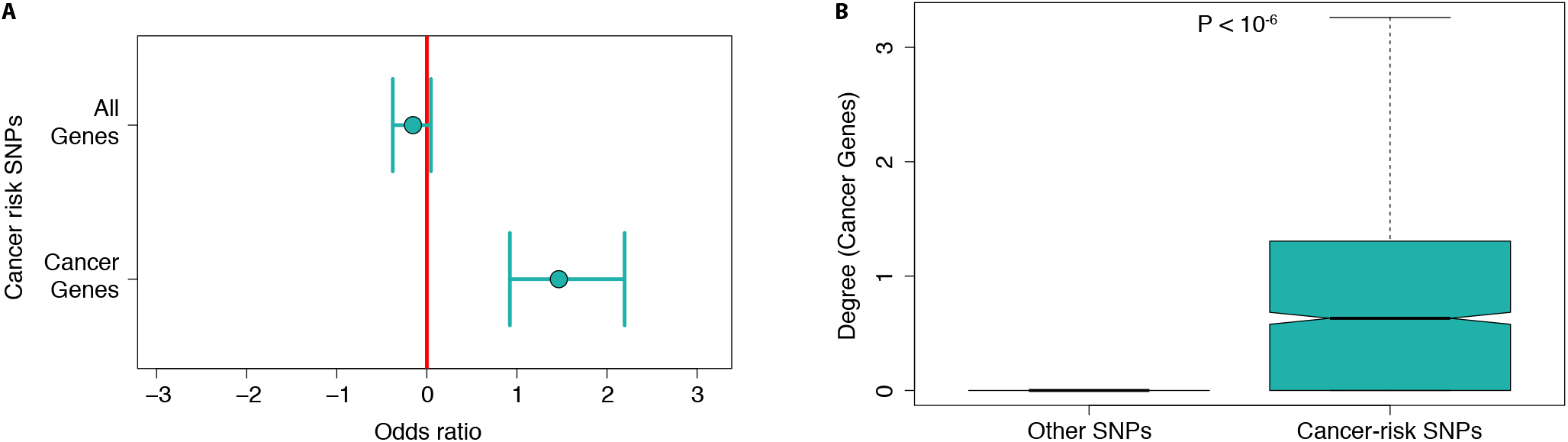
Cancer-risk SNPs are located preferentially in promoters of cancer genes. **A.** Cancer-risk SNPs are preferentially located in the promoters of oncogenes and tumor suppressor genes relative to other genes. This figure shows the odds ratio for finding cancer-risk SNPs, rather than other SNPs, in promoters of all genes’ promoters (top) or oncogenes and tumor suppressor genes’ promoters (bottom). The same analysis for each tissue-specific eQTL network is presented in Figure S3. **B.** Cancer risk SNPs preferentially target oncogenes and tumor suppressor genes across all tissues. Box plots present distributions of number of tumor suppressor genes and oncogenes targeted by cancer-risk SNPs and other SNPs. The P value was obtained using 10^6^ resamplings, taking into account global differences in degree distribution between cancer-risk SNPs and other SNPs. This indicates that cancer genes are likely associated with one or more cancer-risk SNPs, but not other eQTL SNPs. The same analysis for each tissue-specific network is presented in Figure S1.

Finally, we tested whether the cancer-risk SNPs are located in the promoters of genes known to be mutated in cancers. For genes expressed in at least one of the thirteen tissues, we mapped SNPs with minor allele frequencies greater than 5% to their promoters. We then compared SNPs mapping to cancer and non-cancer genes. We found that cancer-risk SNPs are not more likely than other SNPs to fall in promoter regions of non-cancer genes, but cancer-risk SNPs appear preferentially in the promoters of oncogenes and tumor suppressor genes (Figure 3A, supplementary Figure S3).

## Discussion

It has long been known that both germline and somatic mutations in oncogenes and tumor suppressor genes drive development and progression of cancer[?]. However, we know that cancer has a genetic component beyond these well known “cancer drivers,” and that genetic factors can influence differences in the natural history of cancer in individuals possessing the same somatic mutations. Genome-wide association studies have analyzed hundreds of thousands of individuals to find genetic variants that are associated with increased risk of developing cancer, but many of these fall into intergenic regions and have no clear functional association with cancer drivers. As a result, the functional link between genetic risk and the mechanism of cancer development has not been fully understood.

Using data from GTEx, we built bipartite eQTL networks representing germline SNP-gene associations, including both *cis-* and *trans*-acting eQTLs in thirteen different tissues[9]. When we mapped germline cancer-risk SNPs to each of these networks, we found that cancer-risk SNPs are associated with the expression levels of oncogenes and tumor suppressor genes at a far greater rate than expected by chance. This indicates not only that mutations in these cancer genes are important, but also that the genetic control of these genes by regulatory variants plays an important role. A natural assumption might be that cancer-risk SNPs lie in the promoter regions of oncogenes and tumor suppressor genes, but many of the GWAS cancer-risk SNPs are located outside of promoters, leaving the question of the mechanism by which these variants exert their influence.

As we reported previously, SNP-gene eQTL networks are organized into highly modular, regulatory communities that are frequently enriched for genes carrying out distinct biological functions. Consistent with our previous analysis of disease-associated SNPs[8][9], we find that cancer-risk SNPs are overrepresented at the “cores” of individual communities, meaning that those SNPs are at key positions in functional communities where the cancer-risk SNPs can influence the expression of groups of functionally related genes, thus exerting a substantial effect on key biological processes.

Despite the observed concentration of GWAS SNPs in the core of communities, we find that disease-associated germline SNPs in cancer and chronic diseases are distributed differently across eQTL network communities. In chronic obstructive pulmonary disease (COPD), GWAS SNPs map to a small number of communities that possess disease-relevant functions[8]. In contrast, we find that cancer-risk SNPs are distributed across a large number of functionally diverse communities; this distribution is consistent with our understanding that cancer is a systemic disease that affects many different cellular processes.

When we search for communities with the greatest enrichment of cancer-risk SNPs across all thirteen GTEx tissues, we find an over-representation of these SNPs in communities enriched for immune-related genes. In particular, cancer-risk SNPs are linked to altered expression of Major Histocompatibility Complex (MHC) class I and II genes. MHC genes are clustered on the p-arm of chromosome 6 and play a role in recognizing pathogen-infected and other types of modified cells (including cancer cells) and in triggering the innate and adaptive immune system. The high recombination rate and high density of SNPs and genes in the MHC region makes association studies difficult. However, most of the eQTL associations in the region are in *cis,* and some of these have been found in previous studies that targeted the MHC region[20, 21, 22], lending support to our findings. By modulating the expression of MHC genes, cancer-risk SNPs may be modifying an individual’s immune response so as to interfere with the elimination of mutated, pre-cancer cells. Indeed, those eQTL-associated immune response genes belong to the MHC class I and II regions that known to be down-regulated in most cancer cells and affect genes that are targets for some of the newest cancer therapies[23, 24].

In addition to the association with immune response observed in all thirteen tissues, cancer-risk SNPs are over-represented in other functionally interesting communities. For example, SNPs that have been linked in GWASes to breast and epithelial cancer cluster in one eQTL network community in the skin network; a community that is enriched for genes linked to epithelium development and extra-cellular matrix secretion. These and other examples suggest that the distribution of these SNPs within and among communities provides evidence for the functional significance of germline SNPs that are associated with cancer risk and development. It is particularly notable that while the cancer-risk SNPs that associate with gene expression differ between tissues, those diverse SNPs are generally associated through the eQTL network community structure with common functions across tissues. This suggests that similar mechanisms, moderated by tissue-specific expression, may be perturbed across many cancers. This, in turn, may well point to common disease-associated functions that could be targeted therapeutically.

Most importantly, we confirm that the method used here provides an efficient way to explore the effect of germline genetic variants on the risk of cancer. Representing eQTLs using a bipartite network in thirteen tissues, we find that SNPs and genes are organized into communities that reflect the genetic regulatory influence of SNPs on functionally related groups of genes, as demonstrated by GWAS annotation, gene ontology analyses, and enrichment of cancer-risk SNPs in the promoters of cancer genes. By mapping disease-risk SNPs to these networks, we can develop hypotheses about how these SNPs work individually and collectively to moderate risk and possibly enable disease development.

It is well known that the power of eQTL studies to detect associations between genotype and gene expression level depends with the minor allele frequency [25, 26]. In this study, we used data from thirteen tissues for which we had available matching RNA-seq and genotyping data in 200 or more samples; the sample sizes vary between 212 (HRV—heart left ventricle) and 378 samples (SKN—skin). Even the largest sample size does not allow us to reach the maximum power of eQTL detection for alleles with low-intermediate frequencies, and so our results are likely to be enriched for high-intermediate frequency alleles. Because the MHC region is known to include many SNPs with high minor allele frequencies [27], we may be over-estimating its role in cancer risk relatively to other loci. However, the identification of MHC is consistent with the increased recognition of the role that immune processes play in cancer development.

This study provides the first systematic analysis of the regulatory role of germline cancer-risk SNPs and highlights new evidence on the collective regulatory role they play. By mapping cancer-risk SNPs to bipartite networks built from both *cis*- and *trans*-eQTLs in thirteen tissues, we show that cancer-risk SNPs play a particular role in the structure of the eQTL networks, altering the expression of groups of functionally related genes, providing insight into the ways in which these SNPs can increase risk of cancer development. Indeed, this general approach is not limited to cancer, but could be used to provide insight into the functional roles played by other SNPs found to be important through GWASes but for which functional information is not otherwise available. While the approach we implement does not fully bridge the gap between genotype and phenotype, it provides an explanatory framework that can be used to further investigate the genetic risk of disease and the synergistic effects of germline genetic variants.

## Methods

### GTEx data preprocessing, filtering, and merging

We downloaded NHGRI GTEx v 6.0 imputed genotyping data and RNA-seq data (phs000424.v6.p1, 2015-10-05 release) from dbGaP (approved protocol #9112). The RNA-Seq data were preprocessed using Bioconductor `R` YARN package [28, 29] and normalized in a tissue-aware manner using smooth quantile normalization Bioconductor `R` qsmooth package [30]. We identified and removed GTEx-11ILO due to potential sex misannotation. We also filtered out sex-chromosome and mitochondrial genes, retaining 29,242 genes. We excluded five sex-specific tissues (prostate, testis, uterus, vagina, and ovary) and grouped skin samples from the lower leg (sun exposed) and from the suprapubic region (sun unexposed) based on gene expression similarity. For our analysis we only considered tissues for which we had both RNA-seq and imputed genotyping data for at least 200 individuals. Thirteen tissues met all criteria in preprocessing and were used in subsequent analyses (Table S2).

The RNA-seq and genotyping data were mapped by the GTEx Consortium to GENCODE version 19, which was based on human genome build GRCh37.p13 (Sept 2015). We performed principal components analysis on the RNA-Seq data in each tissue, and searched for potentially confounding metadata elements by searching for those correlated with the first ten RNA-Seq principle components. For all tissues, we accounted for the site where the donor was recruited, the RNA extraction kit effects, the quality of extracted RNA, the death place, the time interval between death and start of the tissue sampling, and whether or not the donor was on a ventilator immediately prior to death using the `R` limma package package [31].

### eQTL mapping and bipartite network construction

For eQTL analysis, we excluded SNPs from all analyses if they had a call rate under 0.9 or a minor allele frequency lower than 5% in any tissue. A gene was considered expressed in a sample if its read count was greater than or equal to 6. Genes that were expressed in fewer than 10 of the samples in a tissue were removed for the eQTL analysis in that tissue. To correct for varying degrees of admixture in the African American subjects, we used the first three principal components of the genotyping data provided by the GTEx consortium and included these in our eQTL model.

We used the `R` MatrixEQTL package [7] to calculate eQTLs with an additive linear model that included age, sex and ethnic background, as well as the first three genotype PCs, as covariates:

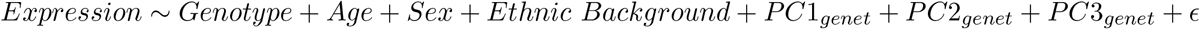

We tested for association between gene expression levels and SNPs both in *cis* and *trans*, where we defined *cis*-SNPs as those within 1MB of the transcription start site of the gene based on mapping using Bioconductor R biomaRt package [32]. P-values were adjusted for multiple testing using Benjamini-Hochberg correction for *cis*- and *trans*-eQTLs separately and only those with adjusted p-values less than 0.2 were used in subsequent analyses.

### Community identification

For each tissue, we represented the significant eQTLs as edges of a bipartite network linking SNPs and gene nodes. To identify highly connected communities of SNPs and genes in the eQTL networks, we used the `R` condor package [8], which maximizes the bipartite modularity [33]. As recursive cluster identification and optimization can be computationally slow, we calculated an initial community structure assignment on the weighted, gene-space projection, using a fast unipartite modularity maximization algorithm [34] available in the `R` igraph package [35], then iteratively converged on a community structure corresponding to a maximum bipartite modularity.

The bipartite modularity is defined in Eq. (1), where *m* is the number of links in the network, *Ã_ij_* is the upper right block of the network adjacency matrix (a binary matrix where a 1 represents a connection between a SNP and a gene and 0 otherwise), *k_i_* is the degree of SNP *i, d_j_* is the degree of gene *j*, and *C_i_*, *C_j_* the community indices of SNP *i* and gene *j*, respectively.

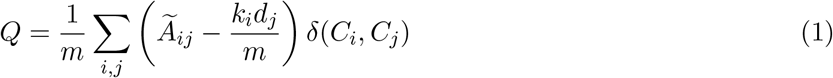

### Cancer risk SNPs

We downloaded the NHGRI-EBI GWAS catalog (Accessed 08 DEC 2015, version v1.0) from the EBI website (https://www.ebi.ac.uk/gwas). We filtered out associations with *P*-values greater than 5 × 10^−8^ and extracted SNPs associated with a risk to develop cancer. We then mapped them to the GTEx data. Specifically, we determined LD blocks using the plink2 –blocks option and a 5MB maximum block size [36] and considered all SNPs in the same LD block as a cancer-risk SNPs as a cancer-risk SNPs.

### Cancer genes

We generated a list of genes mutated in cancers, called ‘cancer genes’ (Supplementary Table S7), including both oncogenes and tumor suppressor genes, by combining information from 2 databases, the Network of Cancer Genes [18] and the Cosmic census [37]. We then mapped the cancer genes to the GTEx data.

We then tested whether the cancer-risk SNPs were preferentially located in the promoters of the cancer genes. We downloaded transcription start site (TSS) positions for all genes present in the GTEx data from the Ensembl database [38, 39] and defined the promoters as the −750/+250bp region around each TSS. We then used a Fisher’s exact test to determine whether the cancer gene promoters were enriched in cancer-risk SNPs. In order to correct for linkage disequilibrium, we used LD blocks rather than SNPs in this analysis.

We tested whether cancer gene SNPs were preferentially targeting cancer genes. To this end, in each network, we computed the SNPs “cancer” degrees for each SNPs, i.e. the number of cancer genes to which each SNP was linked. We then compared the cancer degree distribution between cancer risk and non-cancer-risk SNPs taking into account the global degree distribution using 10^6^ resamplings. We then obtained p-values by comparing the U values obtained from Mann-Whitney U tests on the real and resampled data.

### Cancer risk SNPs enrichments

We tested whether communities were enriched for cancer-risk SNPs using a Fisher’s exact test. We pooled together SNPs from the same LD block and annotated them as cancer risk LD blocks or not cancer risk LD blocks. In each network, we tested whether each community was enriched in each type of cancer risks separately, and then if they were enriched for all cancer risks, taking the whole network as background. To consider a community as enriched in cancer-risk SNPs, we used a threshold of a minimum of 4 LD blocks in the community.

### SNP core score calculation

We defined a SNP’s core score as the SNP’s contribution to the modularity of its community. Specifically, For SNP *i* in community *h*, its core score, *Q_ih_*, is defined by Eq. (2). To normalize SNPs across communities, we accounted for community membership in our downstream testing (Eqns. 3 and 4), which better accounts for community variation compared to the normalization method used in [8].

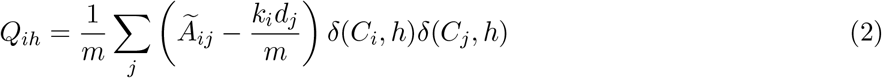

### Gene Ontology functional category enrichment

We extracted the list of genes within each community in each tissue-specific network, and then used the R GOstat package [40] to perform a tissue-by-tissue analysis of the over-representation of Gene Ontology Biological Processes terms within each community enriched for cancer-risk SNPs. Our reference set consisted of all the genes present in the corresponding tissue-specific network. Communities were considered significantly enriched in a given category if the FDR-adjusted p-value was < 0.05.

### Cancer risk SNPs Analysis

We compared the distribution of SNP core scores between cancer-associated SNPs from the NHGRI-EBI catalog and those not associated with traits or diseases for each tissue-specific network using a likelihood ratio test (LRT). In our setting, the LRT assess whether a linear model that includes cancer risk status (Eq. (4)) fits the observed data better than a linear model that doesn’t include this variable (Eq. (3)). As the distribution of SNP core scores (*Q_ih_*) is not uniform across communities, we added community identity as a covariate in the linear regression. In Eqns. 3 and 4, *Q_ih_*, is the core score of SNP *i* in community *h, n* the number of communities in the tissue. *I*(*GW AS* = 1) is an indicator function equal to 1 if the SNP is associated with a higher risk to develop cancer in GWAS and equal to 0 if it’s not associated with any trait or diseases. SNPs associated with traits or diseases other than risk to develop cancer were filtered out. *I*(*C_k_* = 1) is an indicator function equal to 1 if the SNP belongs to community *k* and equal to 0 otherwise.

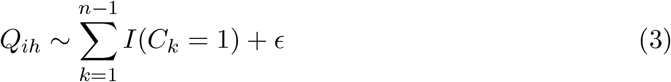

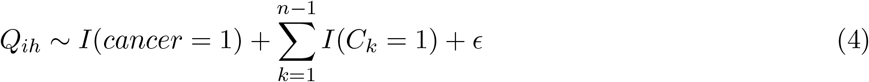

To control for linkage disequilibrium between SNPs, we extracted the median of *Q_ih_* for cancer-risk SNPs and non-GWAS SNPs for each LD block, and used these values as input in the linear regressions.

## Author Contributions

All authors conceived the study; MF, JP and MK analyzed the data; MF, JP, MK, XL and JQ interpreted the results; MF, JP, and JQ drafted the initial manuscript. All authors contributed to the reviewing and editing of the manuscript. All authors read and approved the final manuscript.

## Acknowledgments

This work was supported by grants from the US National Institutes of Health, including grants from the National Heart, Lung, and Blood Institute (5P01HL105339, 5R01HL111759, 5P01HL114501, K25HL133599, K25HL140186), the National Cancer Institute (R35CA220523, 5P50CA127003, 1R35CA197449, 1U01CA190234, 5P30CA006516), and the National Institute of Allergy and Infectious Disease (5R01AI099204). Additional funding was provided through a grant from the NVIDIA foundation. This work was conducted under dbGaP approved protocol #9112.

