## Supplementary Material for "Nongenic cancer-risk SNPs affect oncogenes, tumor suppressor genes, and immune function"

### Supplementary Figures

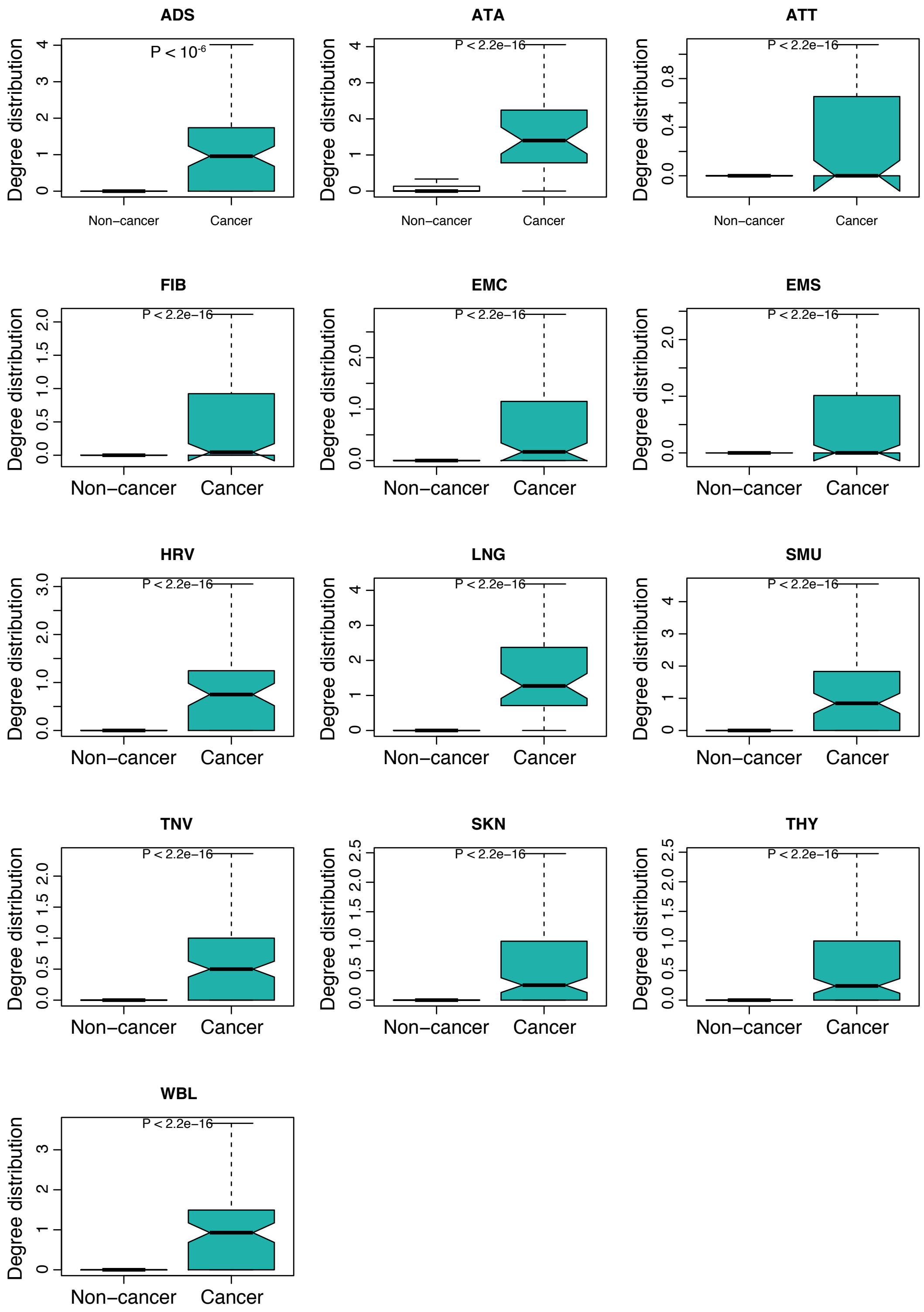


**Figure S1:** Cancer risk SNPs preferentially target oncogenes and tumor suppressor genes. In each panel, box plots present distributions of the number of tumor suppressor genes and oncogenes associated with cancer-risk SNPs and with other SNPs in each tissue-specific network. P values were obtained using $10^6$ resamplings, taking into account global differences in degree distribution between cancer-risk SNPs and other SNPs.

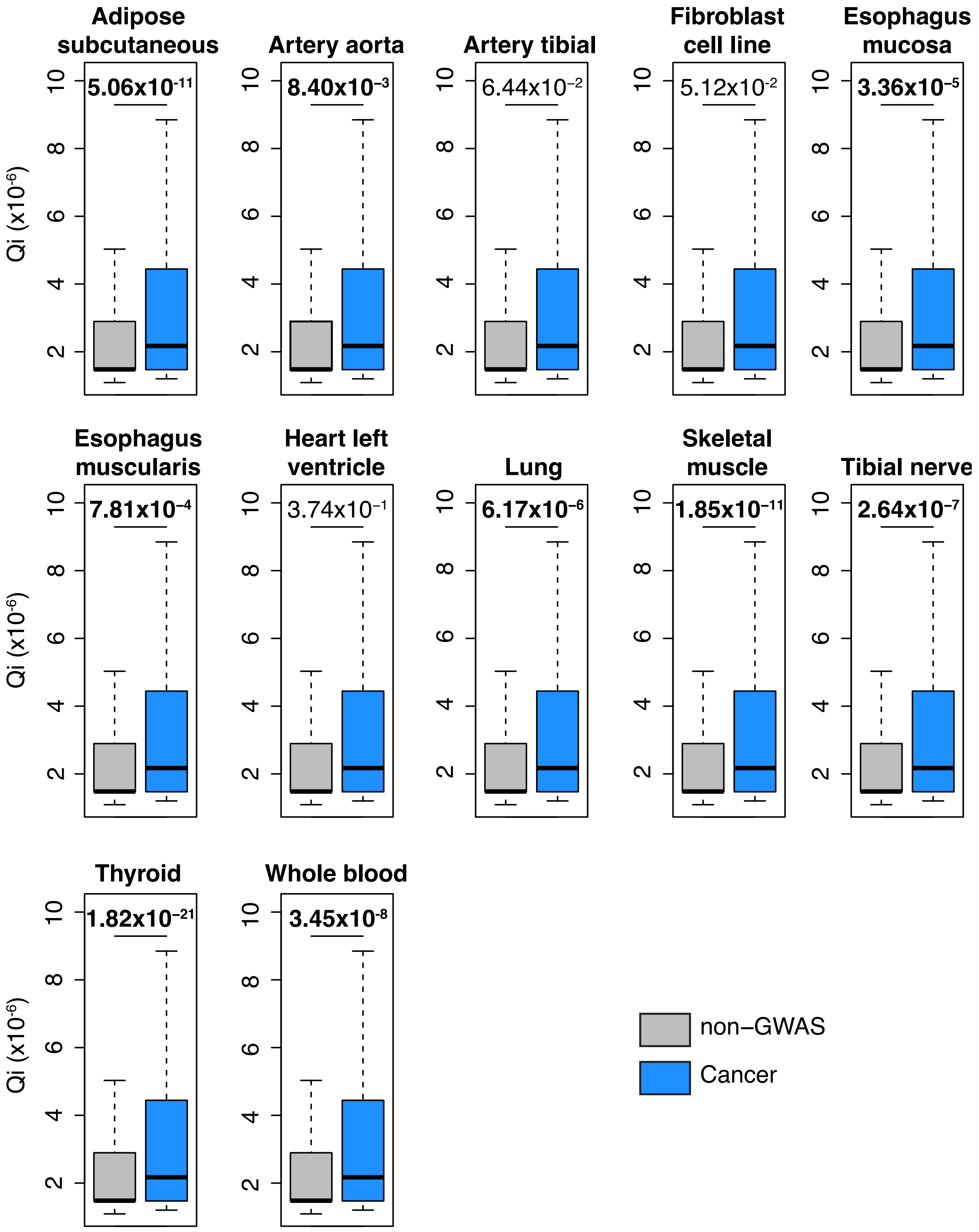


**Figure S2:** Related to Figure 2. SNPs associated in GWAS to a higher risk to develop cancer are enriched in high core score in ten of the thirteen tissues. Significant p-values are indicated in bold text.

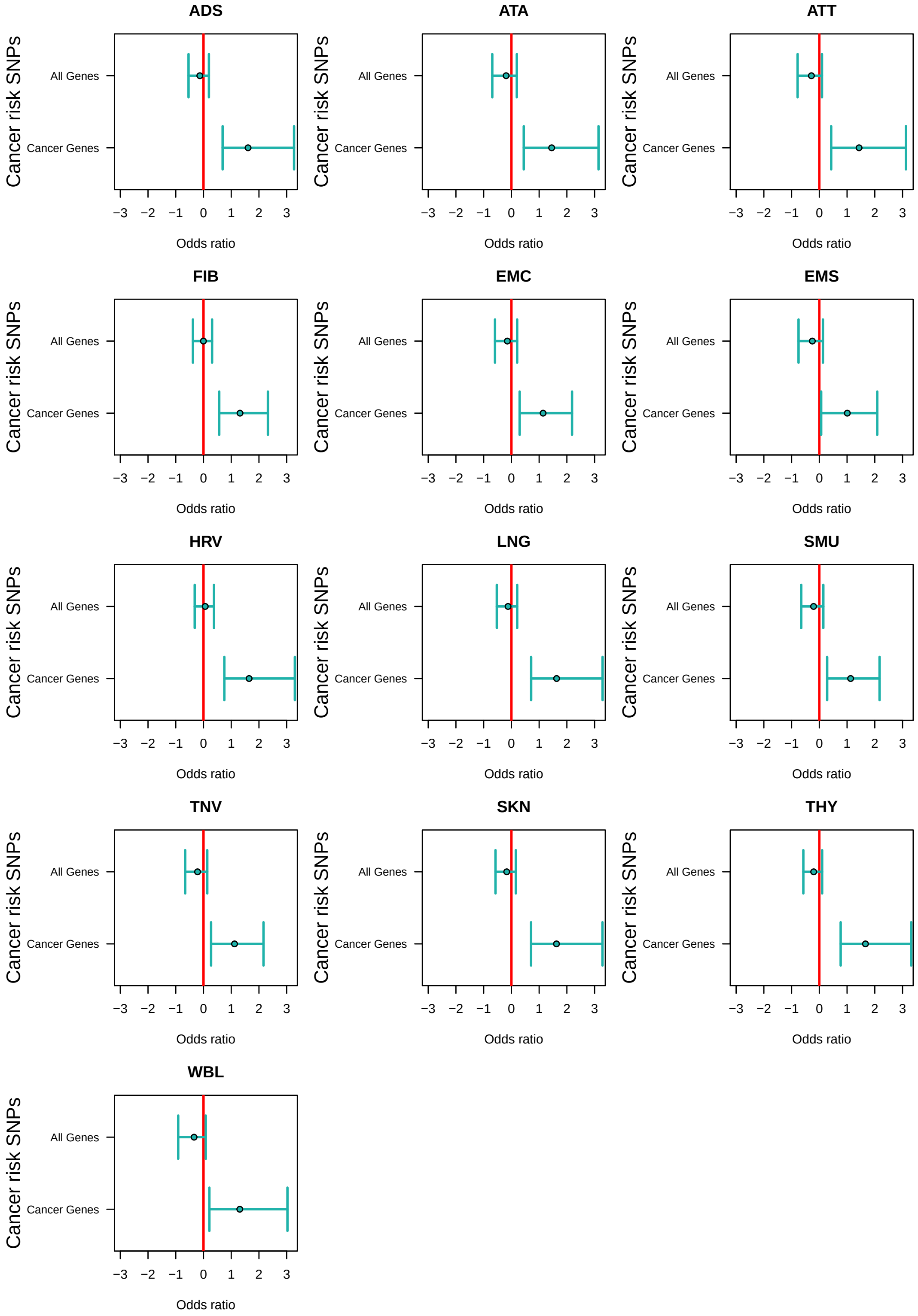


**Figure S3:** Cancer-risk SNPs associated with gene expression levels in eQTL analyses are preferentially located in the promoters of oncogenes and tumor suppressor genes, independent of tissue. The Odds ratio measures the enrichment in cancer-risk SNPs among all genes' promoters (top) or oncogenes and tumor suppressor genes' promoters (bottom) compared to other SNPs.

### Supplementary Tables

**Table S1:** List of cancer-related terms in EBI-GWAS catalog. See Supplementary_Table_1.txt

**Table S2:** Summary of data: tissues and number of samples.

| Tissue names | Abbreviation | GTEx tissues | Samples |
|---|---|---|---|
| Adipose subcutaneous | ADS | Adipose - Subcutaneous | 313 |
| Aorta | ATA | Artery - Aorta | 216 |
| Artery tibial | ATT | Artery - Tibial | 303 |
| Fibroblast | FIB | Skin cells - Transformed fibroblasts | 291 |
| Esophagus mucosa | EMC | Esophagus - Mucosa | 276 |
| Esophagus muscularis | EMS | Esophagus - Muscularis | 245 |
| Heart left ventricle | HRV | Heart - Left ventricle | 212 |
| Lung | LNG | Lung | 290 |
| Skeletal muscle | SMU | Muscle - Skeletal | 378 |
| Tibial nerve | TNV | Nerve - Tibial | 278 |
| | | Skin - Not sun exposed (Suprapubic) | 127 |
| Skin | SKN | Skin - Sun exposed (Lower leg) | 243 |
| | | Total | 370 |
| Thyroid | THY | Thyroid | 295 |
| Whole blood | WBL | Whole blood | 365 |

**Table S3:** Gene Ontology enrichment for communities enriched in cancer-risk SNPs. See Supplementary_Table_4.xlsx

**Table S4:** Cancer-risk SNPs mapping to communities and corresponding Gene Ontology enrichment. See Supplementary_Table_3.xlsx

**Table S5:** Enrichment in cancer-risk SNPs among communities. See Supplementary_Table_5.xlsx

**Table S6:** An eQTL example : genes associated with rs72699833

| Ensembl ID | HGNC | Chr | Start | End | Type | t.stat | P | FDR |
|---|---|---|---|---|---|---|---|---|
| ENSG00000092621.7 | **PHGDH** | 1 | 120,202,421 | 120,286,838 | cis | -3.80 | 1.67E-04 | 2.97E-02 |
| ENSG00000159166.9 | LAD1 | 1 | 201,342,372 | 201,368,736 | trans | -5.62 | 3.73E-08 | 1.93E-01 |
| ENSG00000065618.12 | COL17A1 | 10 | 105,791,044 | 105,845,760 | trans | -5.61 | 3.97E-08 | 1.99E-01 |
| ENSG00000186395.6 | **KRT10** | 17 | 38,974,369 | 38,978,847 | trans | -6.02 | 4.24E-09 | 4.12E-02 |
| ENSG00000178934.4 | **LGALS7B** | 19 | 39,279,851 | 39,282,389 | trans | -6.00 | 4.84E-09 | 4.60E-02 |
| ENSG00000101311.11 | FERMT1 | 20 | 6,055,492 | 6,104,191 | trans | -5.70 | 2.54E-08 | 1.53E-01 |

**Table S7:** Tumor suppressor genes and Oncogenes IDs. See Supplementary_Table_7.txt
